# Functional assessments of *PTEN* variants using machine-assisted phenotype scoring

**DOI:** 10.1101/2020.10.16.342915

**Authors:** Jesse T. Chao, Calvin D. Roskelley, Christopher J.R. Loewen

## Abstract

Genetic testing is widely used in evaluating a patient’s predisposition for developing a malignancy. In the case of cancer, when a functionally impactful inherited mutation (i.e. genetic variant) is identified in a disease-relevant gene, the patient is at elevated risk of developing a lesion in their lifetime. Unfortunately, as the rate and coverage of genetic testing has accelerated, our ability to make informed assessments regarding the functional status of the variants has lagged. Currently, there are two main strategies for assessing variant functions: *in silico* predictions or *in vitro* testing. The first approach is to build generalist computational prediction software using theoretical parameters such as amino acid conservation as feature inputs. These types of software can classify any variant of any gene. Although versatile, their non-specific design and theoretical assumptions result in different models frequently producing conflicting classifications. The second approach is to develop gene-specific assays. Although each assay is tailored to the physiological function of the gene, this approach requires significant investments. Therefore, there is an urgent need for more practical, streamlined and cost-effective methods. To directly address these issues, we designed a new approach of using alterations in protein subcellular localization as a key indicator of loss of function. Thus, new variants can be rapidly functionalized by using high-content microscopy. To facilitate the analysis of large amounts of image data, we developed a new software, named MAPS for machine-assisted phenotype scoring, that utilizes deep learning (DL) techniques to extract and classify cell-level phenotypes. This new Python-based toolkit helps users leverage commercial cloud-based DL services that are easy to train and deploy to fit varying experimental conditions. Model training is entirely code-free and can be done with limited number of images. Users simply input the trained endpoints into MAPS to accomplish cell detection, phenotype discovery and phenotype classification. Thus, MAPS allows cell biologists to easily apply DL to accelerate their image analysis workflow.

## INTRODUCTION

Extrapolating quantitative data from morphological observations enables rigorous statistical analyses; thus, it is a core objective of most cell biology studies. It is crucial that this step is carried out objectively so that one can interrogate the effects of different experimental conditions or variabilities between samples. Traditionally, quantification is accomplished by analyzing images using pre-determined criteria such as cell size, cell shape or protein translocation. However, recent technological advancements in microscopy, such as those in resolution and automation, have empowered us to capture images in greater detail or with higher throughput. However, these advancements also significantly increase the data burden, and the traditional way of manually adjudicating or measuring cellular and subcellular phenotypes cannot be scaled to keep up with the increasing data load. As a result, demands for automated image analysis strategies have surged.

Computational image analysis techniques, which is a part of the larger interdisciplinary field of computer vision, can be grossly divided into those that utilize machine learning and those that do not. Classical computer vision algorithms are stable and efficient, and are already widely used by cell biologists since many of them are packaged into open software platforms like ImageJ and CellProfiler [1]. On the other hand, machine learning techniques distinguish themselves by using iterative cycles of training and fitting that simulates the human learning process of decision making and are more flexible at adapting to the problem at the expense of training time. For instance, given a task of segmenting cells (i.e. detecting individual cell boundaries) in microscopy images, traditional computer vision techniques include thresholding, edge detection, or watershed, while machine learning techniques include clustering, artificial neural network, random forest or support vector machine [2]. Virtually all methods can achieve good performance if they are well-suited and finely tuned for the task. Also, different computer vision techniques are frequently used in concert when building up an image analysis pipeline.

Machine learning, in particular, is an attractive solution for classifying image-based phenotypes due to its ability to extrapolate patterns in the data and make predictions. This approach has been used to screen cell size mutants and to screen small-molecule therapeutics [3,4]. The success of machine learning models requires careful feature engineering in which quantitative measures such as cell shape, pixel intensity and texture are derived from single-cell image data [5]. However, these features need to be predefined, and the initial high-dimensional feature space will require feature selection and feature reduction before it can be effectively used to train the machine learning classifier [6]. Thus, this type of analysis pipeline is usually hand-tuned for each dataset and cannot easily incorporate new data or be transferred to a different dataset.

To overcome this challenge, a specialized branch of machine learning, deep learning, has recently gained favor in the computer vision field. Deep learning is based on the use of multiple layers artificial neural networks and does not require features to be predefined. In computer vision applications, a series of convolutional filters are designed into the network to extract features from pixel-level data to train a deep neural network (i.e. convolutional neural network (CNN)) [7]. This learning structure is inherently flexible at handling a wide variety of image data, and trained networks can also be updated with new data through transfer learning [8]. Thus, CNNs have accelerated computer vision research because of their ability to solve challenging image processing problems, such as 2D/ 3D cell segmentation, organelle segmentation, cell detection and even false fluorescent labeling [9–12]. Further, deep learning techniques are well-suited for automating the analysis of high-throughput (HTP) microscopy data. For instance, CNNs have been used to classify the localizations of fluorescently-tagged proteins in yeast from HTP microscopy images and achieved higher accuracy than traditional machine learning methods [10,13].

However, building deep learning models usually requires substantial programming knowledge that is beyond the means of most biologists. Here, we present a pipeline of using high-throughput microscopy to rapidly detect changes in protein localisations caused by genetic variations. Using the tumor suppressor *PTEN* as proof-of-concept, we show that alterations in PTEN subcellular localization correlated with the pathogenicity of its missense mutations (ie. variants). We also demonstrated a custom-built automated image analysis tool kit that we call MAPS (machine-assisted phenotype scoring) which will help cell biologists leverage the power of deep learning without the need to write any code for model building.

## RESULTS AND DISCUSSION

Many efforts have been devoted to the classification of cancer-related genetic variants. One approach is to develop assays that interrogate the biochemical function of the protein product, and quantitatively measure how variants affect such function. This gene-specific approach has been applied to *BRCA1* variants, where the homology-directed DNA repair function of BRCA1 is the key measure [14]; to *EGFR* variants, where the transforming potential of EGFR is used to assess its mutants [15]; and to *TP53*, where the anti-proliferative function of p53 is used to annotate its variants [16]. Alternatively, another approach is to develop gene-agnostic, generalizable assays by exploiting universal attributes of gene products. One technique measures the relative intracellular abundance of variants which correlate with their pathogenicity [17]. Another technique uses gene expression profiling to fingerprint the molecular functions of a gene and to reveal changes induced by its variants [18].

Previously, we developed a gene-specific assay for the tumor suppressor gene *PTEN* [19]. Although the assay is clinically relevant and scalable, we wanted to engineer a universal multiplexing assay that can be used to functionally assess potentially any gene without needing prior knowledge of gene function. It is well-recognized that the subcellular localizations of proteins are usually crucial for their functions. For instance, the DNA repair activities of p53 and BRCA1 are dependent on their localizations to the nucleus and mutations that disrupt their localizations will significantly impede their functions [20]. Thus, screening for mutations that alter the wildtype protein’s localization could potentially help discover evidence of pathogenicity.

We first established the workflow for scoring the localizations of *PTEN* variants (Figure 1A). We cloned different *PTEN* alleles into an expression vector that expresses GFP and PTEN as a fusion protein interspersed by a P2A self-cleaving peptide, the same design as we previously published [19]. GFP and PTEN would then be expressed as individually folded proteins in 1:1 ratio. After transfection, we carried out immunofluorescence (IF) to visualize PTEN localizations. Finally, we used high-content microscopy to capture images and our custom software to perform automated image analysis and phenotype scoring.

**Figure 1.**
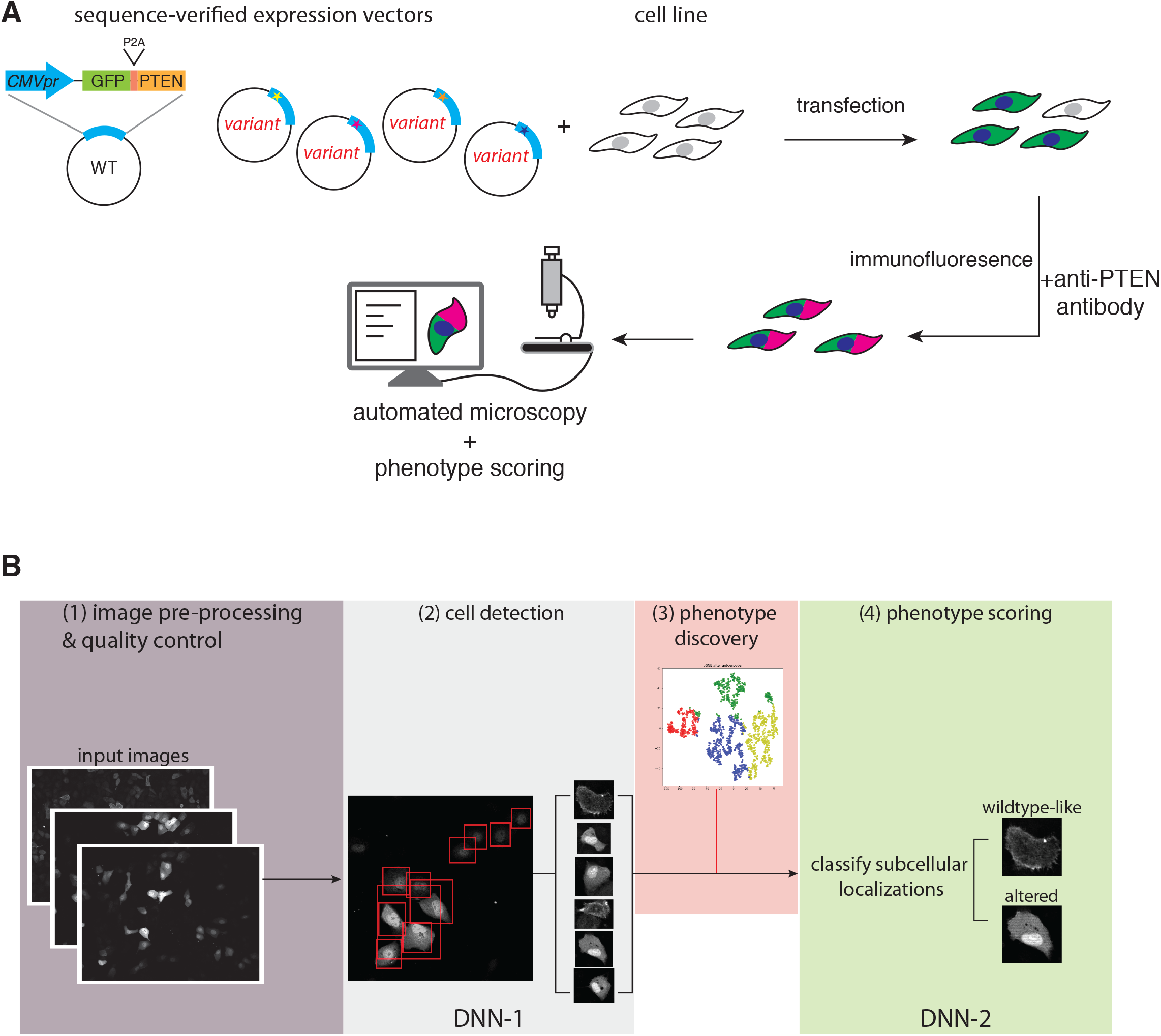
(A) Workflow for expressing and visualizing variants via immunofluorescence and high-content microscopy. (B) Workflow for the MAPS software: (1) Images acquired by high-content microscopy are first pre-processed for quality control (see Figure 3). (2) The first deep neuro network (DNN-1) detects individual cells, giving rise to an initial cell collection (see Figure 4). (3) Feature extraction and 2D manifold embedding help identify unique phenotypes and eliminate outliers (see Figure 5). (4) DNN-2 performs automated phenotype scoring (see Figure 6).

With the goal of streamlining automated phenotype scoring, we developed MAPS, which was designed to score variant localizations, but we anticipate it to have wider applications. MAPS has the following modules: (1) image pre-processing, (2) cell detection, (3) feature extraction/ phenotype discovery and (4) phenotype scoring (Figure 1B). Each of these modules can be executed independently, allowing users to incorporate or substitute their preferred software, such as specialized programs like CellProfiler or image analysis packages like MATLAB or OpenCV. The source codes of MAPS are clearly laid out in a series of Jupyter Notebooks, and each module is organized into their own Notebook, similar to the design principal of the Allen Cell Structure Segmenter [21]. To give a brief overview, the first deep learning model will perform cell detection and isolate individual cells. After, we implemented manifold projection and clustering tools to help the user discover the different classes of phenotypes. Once the user defines the necessary phenotype classes, a final deep learning model will classify all cells identified by the cell detection module.

We carried out pilot experiments to test the pipeline. We first localized PTEN in the non-tumorigenic human breast epithelia cell line MCF10A via IF. Wildtype PTEN has been reported to shuttle between the cytoplasm and the plasma membrane, which is essential for its tumor suppressor function in dephosphorylating phosphatidylinositol-3,4,5 trisphosphate [22]. Although PTEN does not contain canonical nuclear localization signals (NLS), nuclear PTEN is apparent in quiescent cells, but not normally found in dividing cells [23]. Consistently, we noticed that wildtype PTEN localized mostly in the cytosol with minor PM staining, but is excluded from the nucleus. In contrast, the localization of a known tumour-associated loss of function variant, C124R [24,25], was predominantly nuclear (Figure 2). Since there were clear differences in the localizations of selected PTEN alleles, we felt confident to use alterations in PTEN localizations as the key phenotypic measure for scoring variant function.

**Figure 2.**
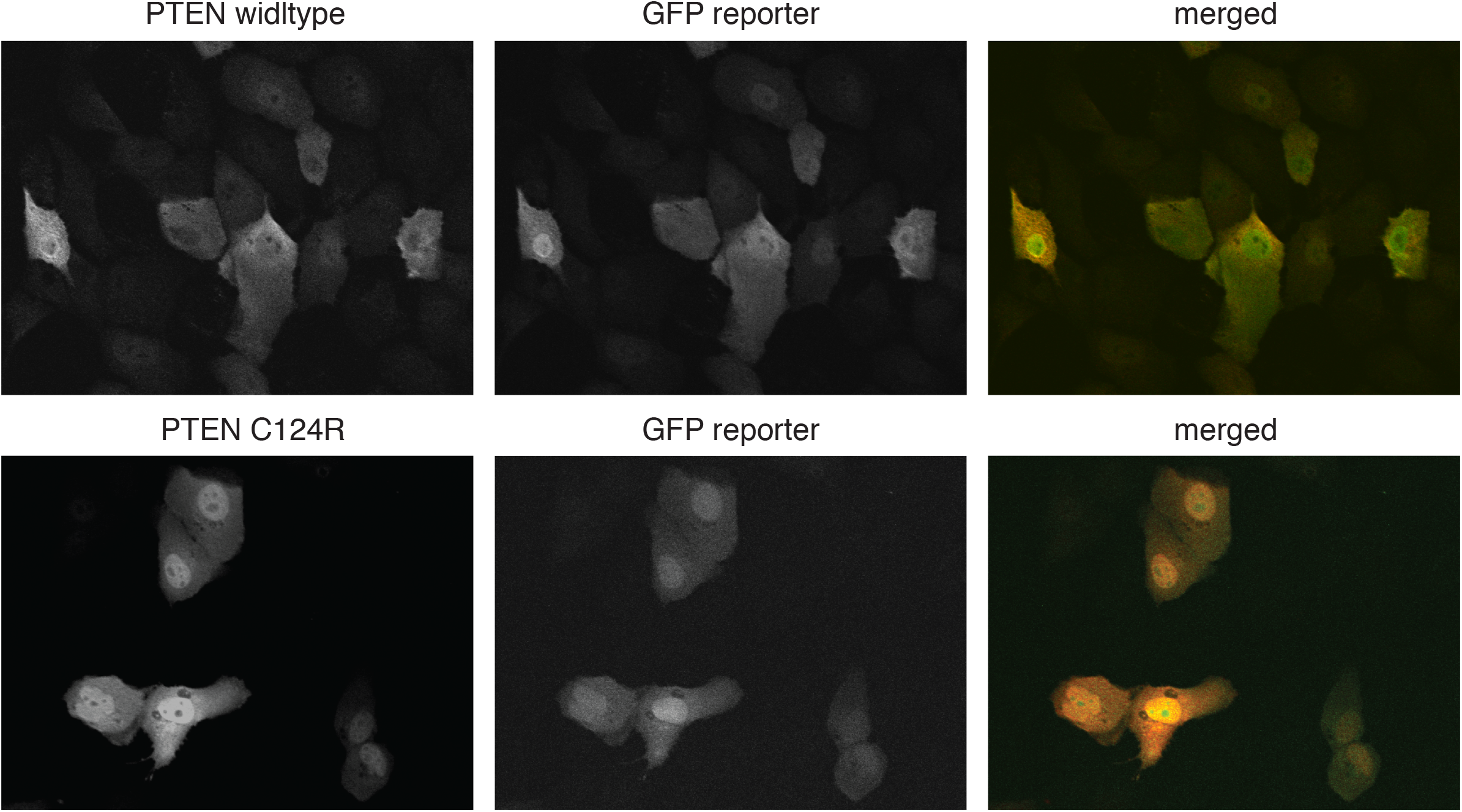
Representative images of the localizations of wildtype and C124R allelic variant of PTEN as visualized via immunofluorescence. PTEN and the GFP reporter were expressed in 1:1 ratio.

Next, we acquired images using HTP microscopy. We expected that automated microscopy instruments and experiments will inevitably generate sub-optimal images such as those that are out of focus or those that contain artifacts including air bubbles, scratches or foreign fibers may be present. Images with these aberrations will need to be removed to maintain the quality of downstream analyses. Thus, after automated image acquisition, a pre-processing step is necessary to perform quality control (QC). Conventional image analysis pipelines would expect users to remove these problematic images prior to analysis. However, automated microscopes can easily generate gigabytes of image data, making manually screening and excluding problematic images time consuming and undesirable. A number of strategies are available to perform image QC, such as building custom software solutions [26] or using the QC pipeline implemented in CellProfiler’s MeasureImageQuality module [27]. In order to integrate seamlessly with the other modules, we implemented custom functions to compute focus measures using OpenCV. We used Variance of Laplacian as the focus measure which measures the amount of edges in an image, so that an in-focus images will have high variance, and a blurry image will have low variance (Figure 3A) [28,29]. However, images containing artifacts such as air bubbles or overexposed cells (due to cells overexpressing the target protein) will create high amount of edges and be mis-labeled as in-focus images. To overcome these issues, we implemented image dilation to remove the edges from air bubbles (Figure 3B), and masking followed by Gaussian blurring to remove edges from overexposed cells (Figure 3C). After screening and mitigating these artifacts that affected focus measure calculations, we achieved on average over 90% accuracy in removing out of focus images.

**Figure 3.**
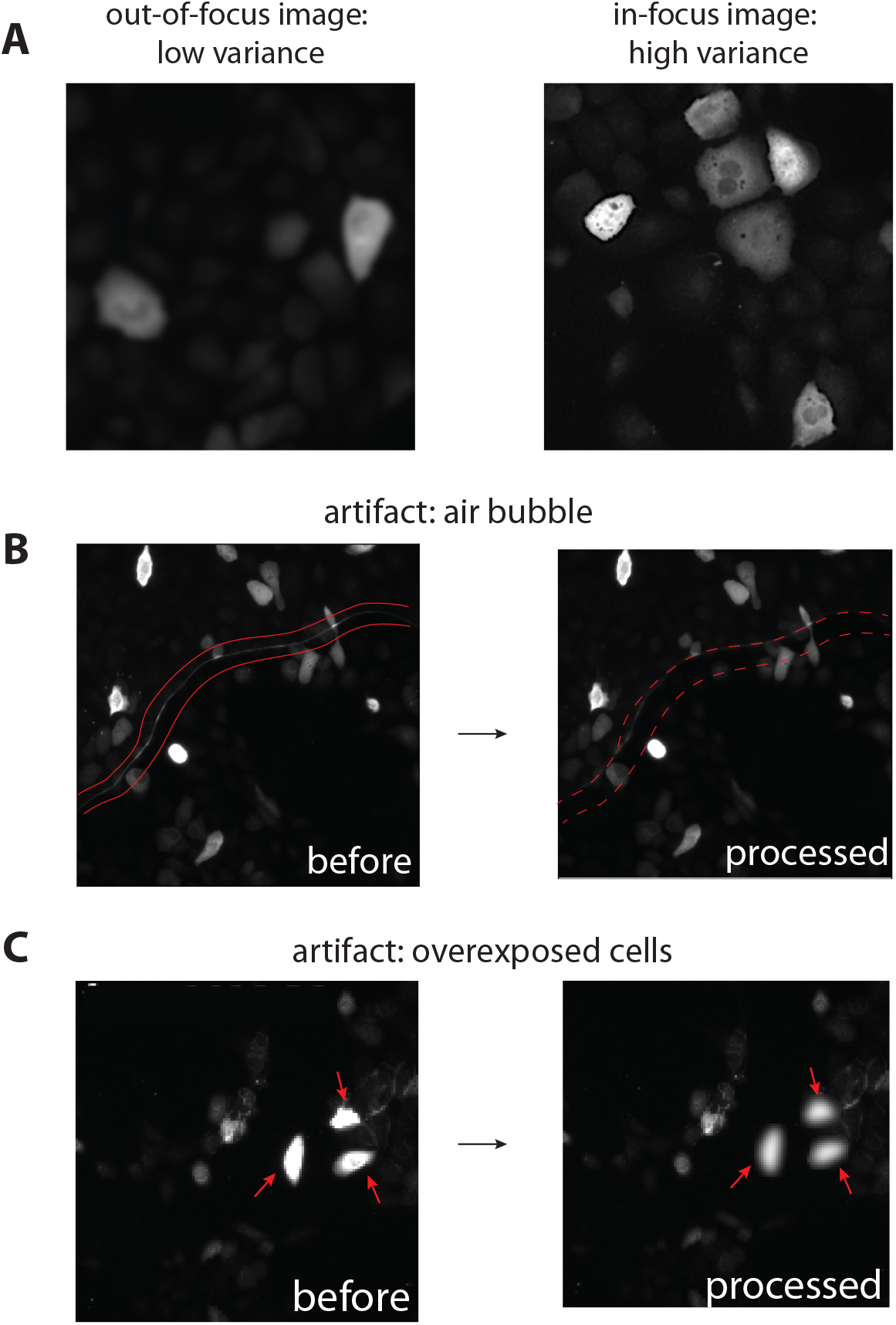
Functional details of image pre-processing and quality control. (A) Laplacian variance is used as a focus measure operator to differentiate blurry and in-focus images. (B and C) Artifacts such as air bubbles or overexposed cells interfere with focus measure calculations and are further processed to obtain accurate focus measures.

After image QC, we need to isolate individual cells to extract cell-level features. Cell detection places bounding boxes around each cell object, and is different from cell segmentation which aims to identify cell boundaries as defined by the plasma membrane (e.g. in mammalian cells) or by the cell wall (e.g. in yeast cells). We realized that in certain cases cell segmentation would be desirable, but cell detection was well-suited for our application. To carry out cell detection, we took advantage of the Custom Vision module of Azure, Microsoft’s cloud-based machine learning platform. We trained a custom cell detection model on Azure and used the endpoint to predict bounding boxes on all of our HTP microscopy images. To improve the performance of the Azure model, we implemented augmentation strategies, a common technique used in CNN training [9], to boost the training data. These included image rotation, flipping, contrast adjustments, color inversions and adding noise (Figure 4A). We boosted training data from 148 to 2200 images, and achieved 9.8% improvement in precision and 3.4% in recall (Figure 4B).

**Figure 4.**
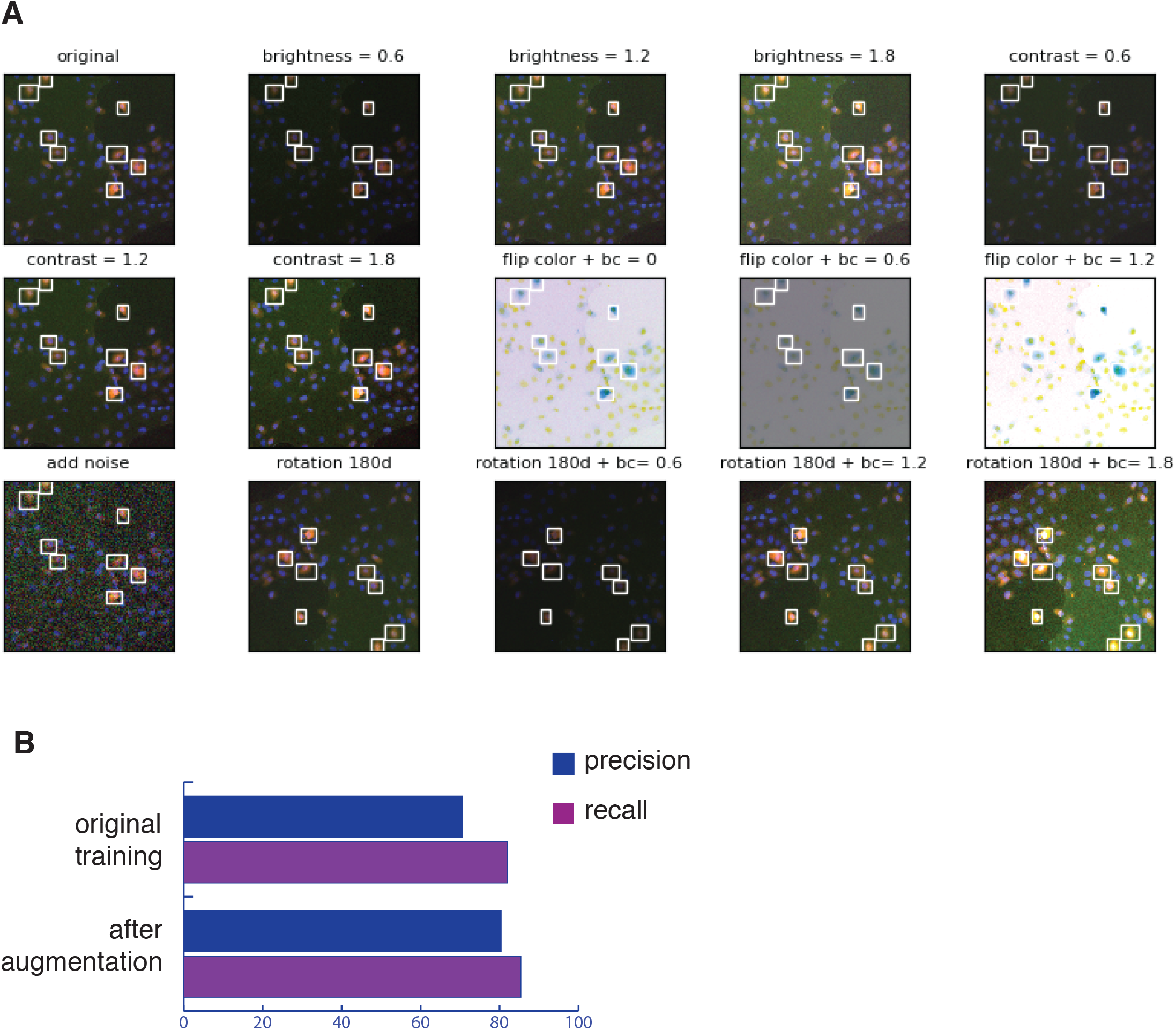
(A) Sample images from training augmentation. The original image (top left) underwent different brightness and contrast (bc) adjustments, color inversion (flip color), noise addition and rotation. (B) The precision and recall performances of the Azure object detection model before and after training augmentation.

Perhaps the most noteworthy distinction between our image analysis pipeline and others is that we let users build experiment-specific deep learning models using commercial solutions. Since imaging experimental conditions vary greatly, it is more practical for users to train and deploy their specific cell detection and phenotype scoring models instead of providing pre-trained models in order to achieve the best accuracy. Also, because deep learning is computation heavy, and users typically have different computer hardware with varying capabilities, we considered using cloud computing platforms will provide the most consistent training and prediction performance at a fraction of the cost of purchasing new hardware. After testing the leading cloud machine learning platform, including Microsoft Azure, Amazon AWS and IBM Watson, Azure has the most intuitive graphical user interface. Importantly, the Custom Vision module on Azure allows users to create computer vision models entirely code free, and can achieve reasonable performance using very few training images (as little as 20 images). This is immensely more convenient than building deep learning models from scratch because doing so typically require hundreds if not thousands of curated images for training and validation [7].

One common approach of phenotype classification is to first quantitatively measure cell morphologies followed by training a ML classifier using the quantified measurements as features [5,30]. It is necessary then to explicitly define these cell-level features, such as cell size, cell shape or pixel intensity [5]. This approach was the standard practice before deep learning became practical and is still widely used today. Alternatively, a different approach is to carry out feature extraction using a convolutional network, which replaces handcrafting features [11]. This is the preferred solution because the same CNN structure can be used to analyze a wide variety of dataset and does not require expert knowledge in designing the types of features to extract. Since we aimed to maximize flexibility, we chose the second approach to only broadly define PTEN’s localization pattern and use CNN to carry out the classification.

The next step in the pipeline was to discover all relevant phenotypes that we planned to train the final classifier. Specifically, we needed to carefully inspect the cropped cells for any unexpected phenotype or outliers. We also should remove low quality images such as objects that were mis-identified as cells. Thus, we implemented useful functions to facilitate visual inspections by clustering cells that show similar morphologies and by generating cell galleries. We designed two different methods to extract cell-level features. The first method involves applying convolutional filters to the input image, followed by parallel stacking of the convolved layers. Then, all layers are flattened to generate a high-dimensional dataframe (Figure 5A). Finally, we used t-SNE to generate a 2D manifold projection of all the cells, and randomly inserted the original cell images to facilitate manual inspection (Figure 5B). We also applied spectral clustering to group cells with similar morphologies. The combination of these techniques helped reveal that the red cluster consisted of junk data from inaccurate cell detection, and cells in the green cluster were too small to be useful for phenotype scoring (Figure 4B). Consequently, we eliminated images from the red and green clusters. The second method is by using a deep autoencoder (Figure 5C). The autoencoder has a contracting path that is very useful for dimensionality reduction and feature learning [31]. After using the autoencoder to perform feature extraction, we similarly generated a t-SNE projection followed by spectral clustering and embedding a cell gallery (Figure 5d). In this example, junk data was grouped into the green cluster, and small cells were in the red cluster. Therefore, either approach can help us quickly remove potentially confounding data, allowing us to focus on inspecting a more manageable subset of images.

**Figure 5.**
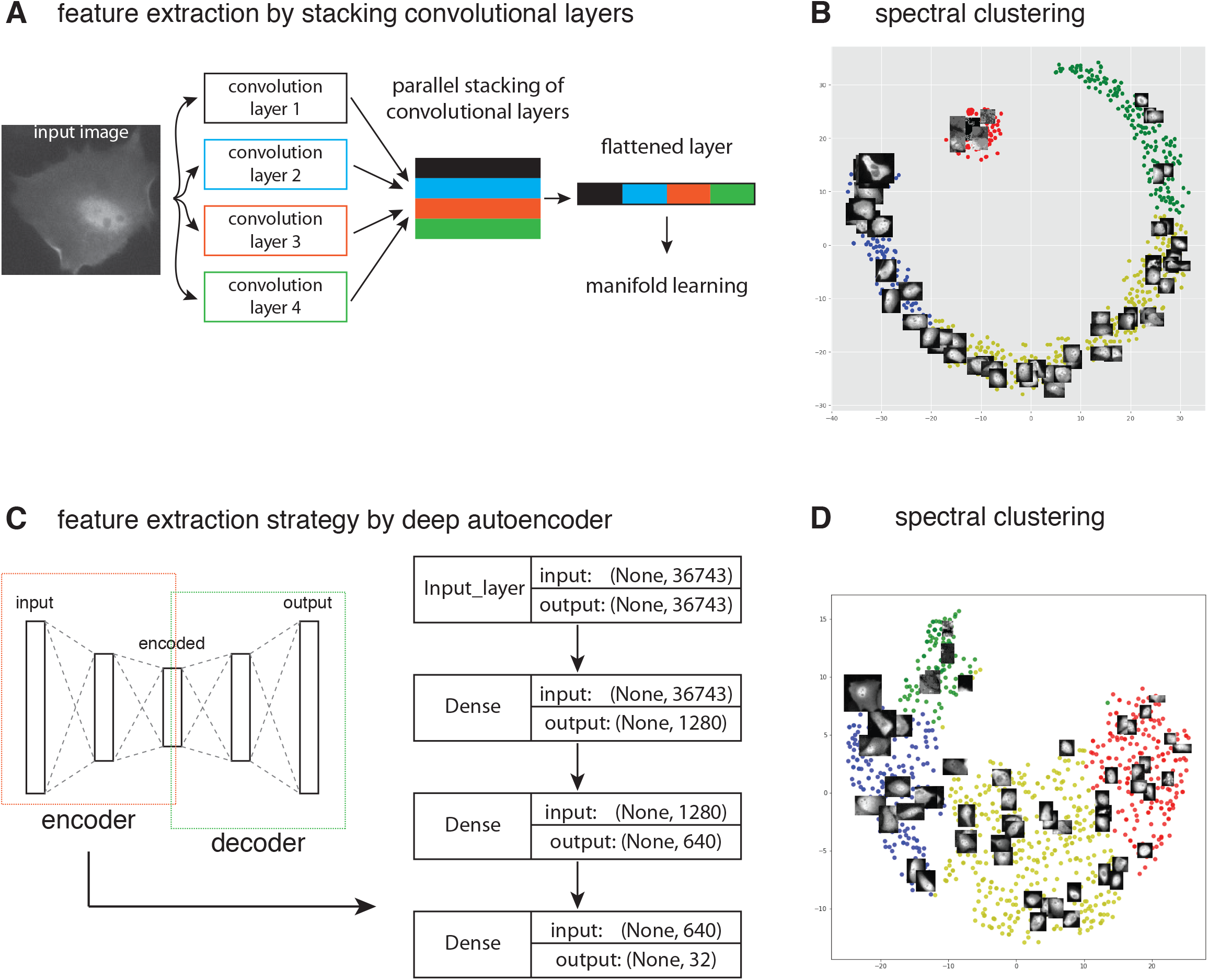
Feature extraction and manifold projection techniques to assist phenotype discovery. (A) Workflow for feature extraction by parallelly stacking convolutional layers followed by flattening them. (B) The flattened layer then underwent dimensionality reduction by t-SNE into 2 components and projected onto a 2D scatter plot. Spectral clustering was applied to find the decision boundaries, and original images of the inputs were also plotted onto the scatter plot. (C) Workflow for feature learning by using a deep autoencoder. Left, schematics of an autoencoder, but only the encoder portion was used to find the latent representation of the input. Right, the encoder architecture. (D) Similar to (B), but the encoded latent representations were the inputs of t-SNE.

After inspecting the remaining images, we discovered that there were three main patterns of PTEN localizations common to all tested variants: non-nuclear, nuclear and diffused (Figure 6A). We trained another classification model on Azure, and plug the endpoint into our script to perform automated phenotype scoring based on these three classes. We analyzed wildtype PTEN and 7 variants: G44D, C124R, G127R, M134I, R173H, Y180H and P246L (Figure 6B). Using our imaging pipeline and MAPS, we were able to analyze large number of cells (n= 350-700). Importantly, we noticed that very few wildtype cells had nuclear PTEN localization, whereas the non-functional variant C124R had significantly higher nuclear PTEN (10% vs 52%), consistent with our initial observation (Figure 2). We also looked up the loss of function (LOF) scores that we previously measured for these variants using a spheroid assay (Figure 6C, [19]). Notably, we found that the percentage of cells with nucleus-localized PTEN correlated strongly with LOF scores (Figure 6D, Pearson’s correlation = 0.737). There were two outliers: R173H had a low LOF score but high nuclear PTEN; G127R had the highest LOF score in our test, but its nuclear PTEN was similar to that of C124R or M134I. We suspected that assessing variant function by subcellular localization may be more sensitive than the spheroid assay, but the spheroid assay can detect a higher degree of LOF. More testing might be able to further dissect the pros and cons of either approaches.

**Figure 6.**
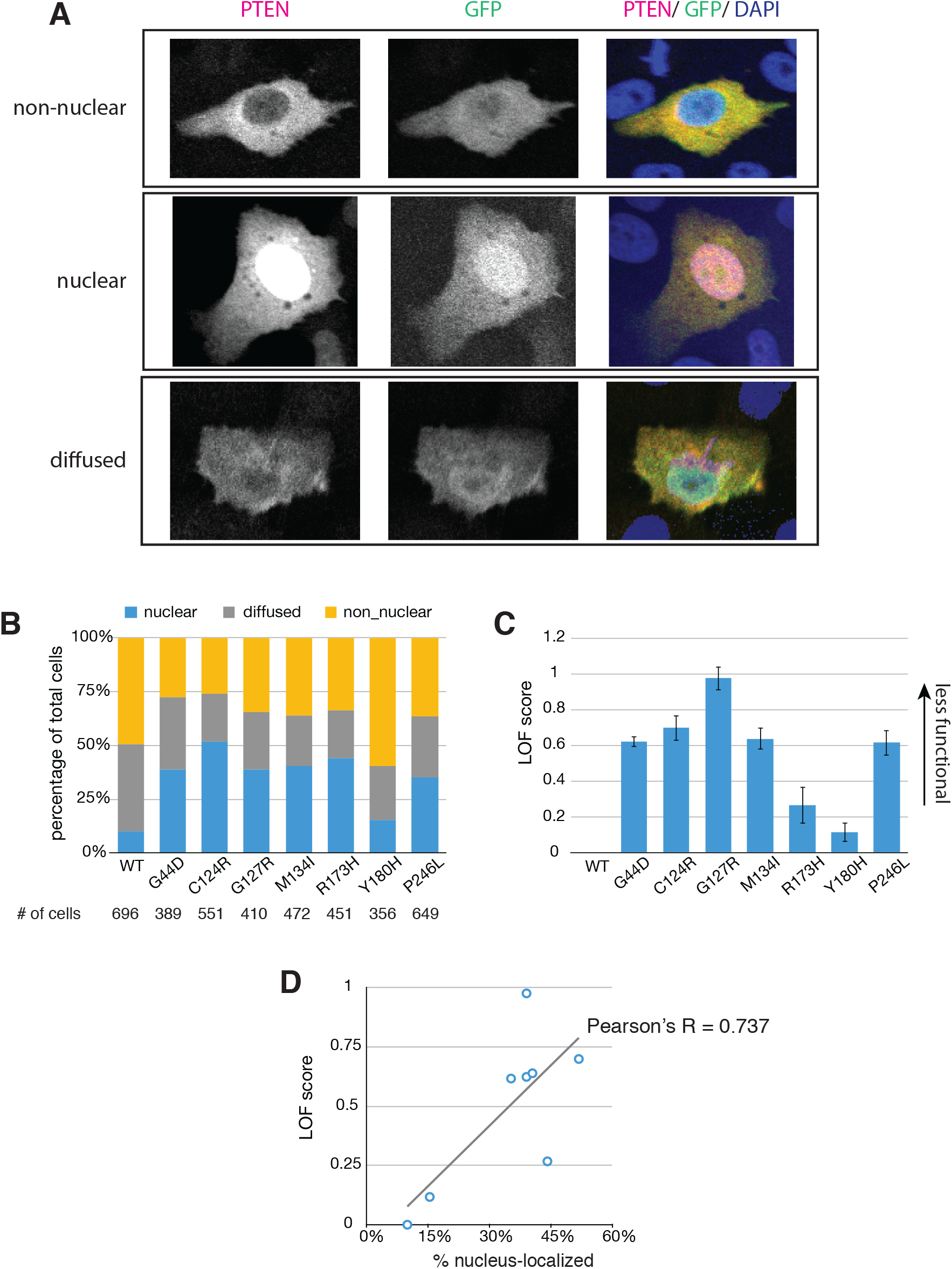
(A) Representative images of the three PTEN localization classes that were used to train the final classification model on Azure. (B) Results of automated phenotype scoring for wildtype PTEN and seven variants. (C) LOF scores for the same variants as in (B) taken from a previously published spheroid assay. (D) Scatter plot showing the correlations between the LOF scores and the percentage of cells with nuclear PTEN for the seven tested variants. The best fit line is also shown.

In conclusion, we developed MAPS specifically to automate phenotype scoring of high-content microscopy data, and used it to score the localizations of PTEN variants. MAPS stands out for other software tools in that it allows users to build custom deep learning object identification and classification models using Microsoft’s Azure cloud computing platform, completely code-free. We hope MAPS can help empower cell biologists with the power of deep learning even if they do not have the expertise in AI. Finally, assessing variant function using high-content microscopy is a simple and easy to scale approach, and could be more cost-effective than developing gene-specific assays.

## MATERIALS AND METHODS

### Cell Culture

The *PTEN*−/− cell line (MCF10A background) was purchased from Horizon Discovery and verified by western blotting. Cell were cultured according to published protocols [32] and were maintained in a 37°C incubator with 5% CO2. Mycoplasma was tested monthly by direct DNA staining with DAPI.

### Plasmids and Transfections

PTEN expression vectors were generated as previously described [19]. Transfection was carried out 24 hours after seeding 50,000 cells in a 12-well dish containing 22×22mm glass coverslips (Thermo Fisher Scientific) using Lipofectamine 2000 (Thermo Fisher Scientific) according to manufacturer’s protocols. Successful transfection was confirmed by direct visualization of GFP expression using a fluorescent microscope.

### Immunofluorescence

24 hours after transfection, cells were fixed using 4% paraformaldehyde in PBS. Cells were permeabilized with 0.1% triton x-100 in PBS, blocked with 10% BSA, and incubated overnight with rabbit PTEN antibody (138G6, Cell Signaling Technology). Coverslips were then incubated with mouse anti-rabbit Alexa Fluor 568-conjugated antibody (Invitrogen), followed by DAPI, and mounted using ProLong Gold antifade mountant (Thermo Fisher Scientific).

### High-content microscopy

Images were acquired using a Cellomics Arrayscan (Cellomics Inc.). using a 20x objective. 500 images were acquired per coverslip at 3 channels (green/ red/ blue) per image.

### MAPS custom software

MAPS was written in Python 3.6.10. Other packages include numpy (1.18.1), pandas (1.0.3), opencv-python (4.1.1.26) and matplotlib (3.1.3). All codes are available at: https://github.com/jessecanada/MAPS.

